# Stimulus-Dependent Theta Rhythmic Activity in Primate V1 Predicts Visual Detection

**DOI:** 10.1101/2021.11.30.470367

**Authors:** P. T. Fanyiwi, B. Agayby, R. Kienitz, M. Haag, J. Cadena-Valencia, M. C. Schmid

## Abstract

Theta-band (3–8 Hz) neural oscillations are integral to sensory processing and active exploration. Traditionally associated with higher-order areas such as hippocampus and prefrontal cortex, recent studies identified theta rhythmic modulations in the primary visual cortex (V1) of mice during locomotion, suggesting sensory processing functions. Here, we demonstrate that careful optimization of visual stimulus size and contrast can induce robust theta oscillations in macaque V1. During visual detection, monkeys’ reaction times fluctuated rhythmically at the theta frequency of V1 neural activity, with detection performance correlated to the theta phase. These findings suggest that induced theta oscillations may reflect an intrinsic temporal filtering mechanism of V1 neurons, highlighting the importance of early sensory cortical dynamics in shaping perceptual timing.

## Introduction

Theta rhythmic neural activity has initially been observed in higher-order cortices, such as the hippocampus and prefrontal cortex (Buzsáki, 2002; Siapas et al., 2005), and is traditionally associated with spatial navigation, reward prediction, and memory. For example, hippocampal theta oscillations, documented during rodent locomotion, are thought to integrate sensory inputs with motor and cognitive processes (Buzsáki, 2002; Vanderwolf, 1969). Recent evidence however reveals theta-band activity in primary sensory areas, like the primary visual cortex (V1) of mice during locomotion and sensory memory formation (Fournier et al., 2020; Niell & Stryker, 2010; Zimmerman et al., 2025), suggesting sensory areas contribute to theta rhythms, linking sensory input with motor behavior and memory. Lesion studies in rodents and monkeys underscore V1’s critical role in generating this sensory theta rhythm (Kienitz et al., 2021; Zimmerman et al., 2025). Despite these insights, the stimulus-driven processes associated with theta oscillations in primary sensory cortex remain poorly understood.

The responses of V1 neurons to visual stimulation are well known to reflect basic stimulus properties such as orientation, spatial frequency, contrast, and size (Carandini et al., 2005). When tested with a drifting grating stimulus, many V1 neurons exhibit preferentially tuned responses to temporal frequencies in the theta range (Foster et al., 1985; Hawken et al., 1996). An interesting hypothesis is therefore that V1 neurons might act as a theta-tuned temporal filter to incoming sensory information, thereby shaping the timing of visual perception and help integrate with theta-dependent processing in higher-order cortices (Fiebelkorn et al., 2018; Fiebelkorn & Kastner, 2021; Helfrich et al., 2018; Jutras et al., 2013) Accordingly, presenting a static visual stimulus could induce theta oscillations consistent with the preferred temporal frequency of V1 neurons and influence the timing of detecting any changes in the stimulus that is consistent with the induced theta rhythm.

To test this hypothesis, we investigated whether theta-rhythmic neural activity in macaque V1 can be externally induced through static visual stimuli varying in size or contrast. As we confirmed that stimulus manipulations could elicit robust theta-band oscillations in V1, we subsequently assessed their contribution to visual detection performance. Our findings extend the understanding of theta-band oscillations in primary sensory areas, showing they can be driven by external stimuli. This is consistent with the hypothesis that V1 neurons temporally filter incoming visual stimuli in theta frequency, which in turn influences perceptual processes.

## Results

A fundamental aspect of V1 neuron responses is that activity strength is modulated by stimulus size, reflecting a spatial summation in the receptive field (RF) (Cavanaugh et al., 2002; Jones et al., 2001). To establish the link between this size tuning of V1 and theta oscillations, we first investigated how stimulus size influences theta-band (3–8 Hz) oscillations in the primary visual cortex (V1) of macaque monkeys. Consistent with established size-tuning properties (Cavanaugh et al., 2002; Jones et al., 2001), increasing stimulus size from 0.3° to 1° elevated neuronal firing rates, while further enlargement to 4° reduced activity (Figure 1A). Notably, a 1° stimulus elicited pronounced theta oscillations, evident in the peri-stimulus time histogram (PSTH) of single-unit activity (SUA), multi-unit activity (MUA), and local field potential (LFP) time courses (Supplementary Figure 1A).

**Figure 1.**
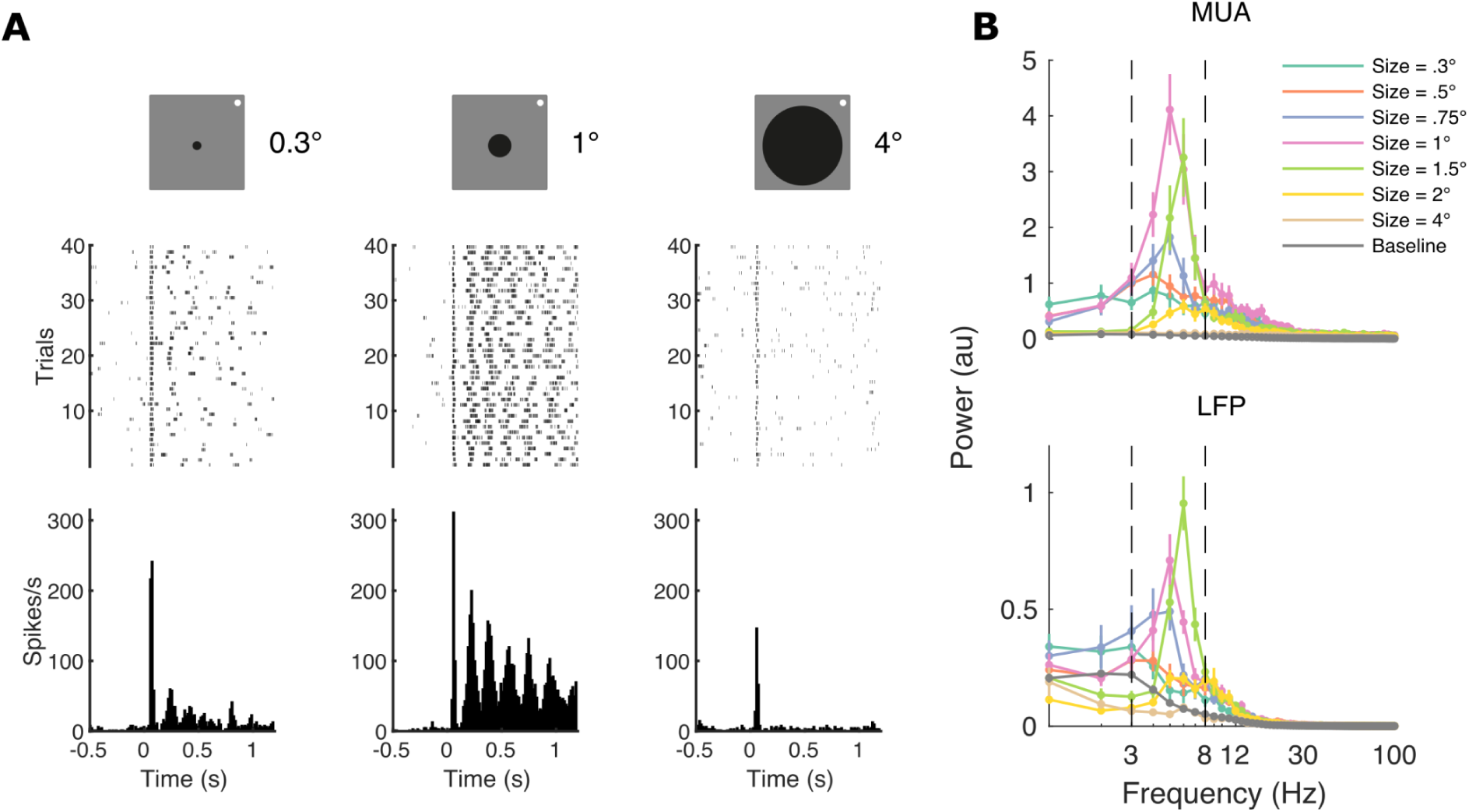
Theta oscillations at different stimulus sizes: example. (A) Top: Schematic of the stimulus, a black disk, presented at three different sizes, with diameter in visual degrees, while the animal (monkey AL) fixated at the white fixation spot. Sizes are not drawn to scale. Middle and bottom: Raster plots and peri stimulus time histogram (PSTH) of an example single unit activity aligned to stimulus onset for 0.3° (left), 1° (middle), and 4° (right). Note the rhythmic activity elicited explicitly by the 1° stimulus. (B) Top: Mean power spectra of multi-unit activity (MUA) for each stimulus size. The MUA was obtained from the same electrode from which the single unit activity in A was isolated. Bottom: Mean power spectra for LFP. Dashed lines delineate theta frequency range. Error bars are ± 1 standard error of the mean (SEM).

To quantify these oscillations, we computed power spectra of the envelope of MUA (Drebitz et al., 2019; Supèr & Roelfsema, 2005) and LFP after stimulus onset (200 ms to 1200 ms post-stimulus) (Figure 1B). We found that specific stimulus sizes consistently induced strong theta rhythmic activity: 1° in MUA and 2° in LFP. These sizes yielded higher theta-band power and more consistent peak frequencies within the theta range across individual trials as compared to other stimulus sizes (Supplementary Figure 1B).

At the population level across channels, theta oscillations in MUA and LFP were significantly modulated by stimulus size in both monkeys (Figure 2A, C; Supplementary Figure 3A) (p < 0.001 for both monkeys; n = 248 for monkey AL MUA, n = 243 for monkey AL LFP, n = 390 for monkey DP MUA and LFP; Friedman test). Both increased with stimulus size up to a point, beyond which further enlargement led to decreases (Figure 2B, D; Supplementary Figure 2). Medium-sized stimuli consistently produced the most substantial theta peaks. Minimal theta activity was observed during pre-stimulus baseline periods and with the smallest and largest stimuli.

**Figure 2.**
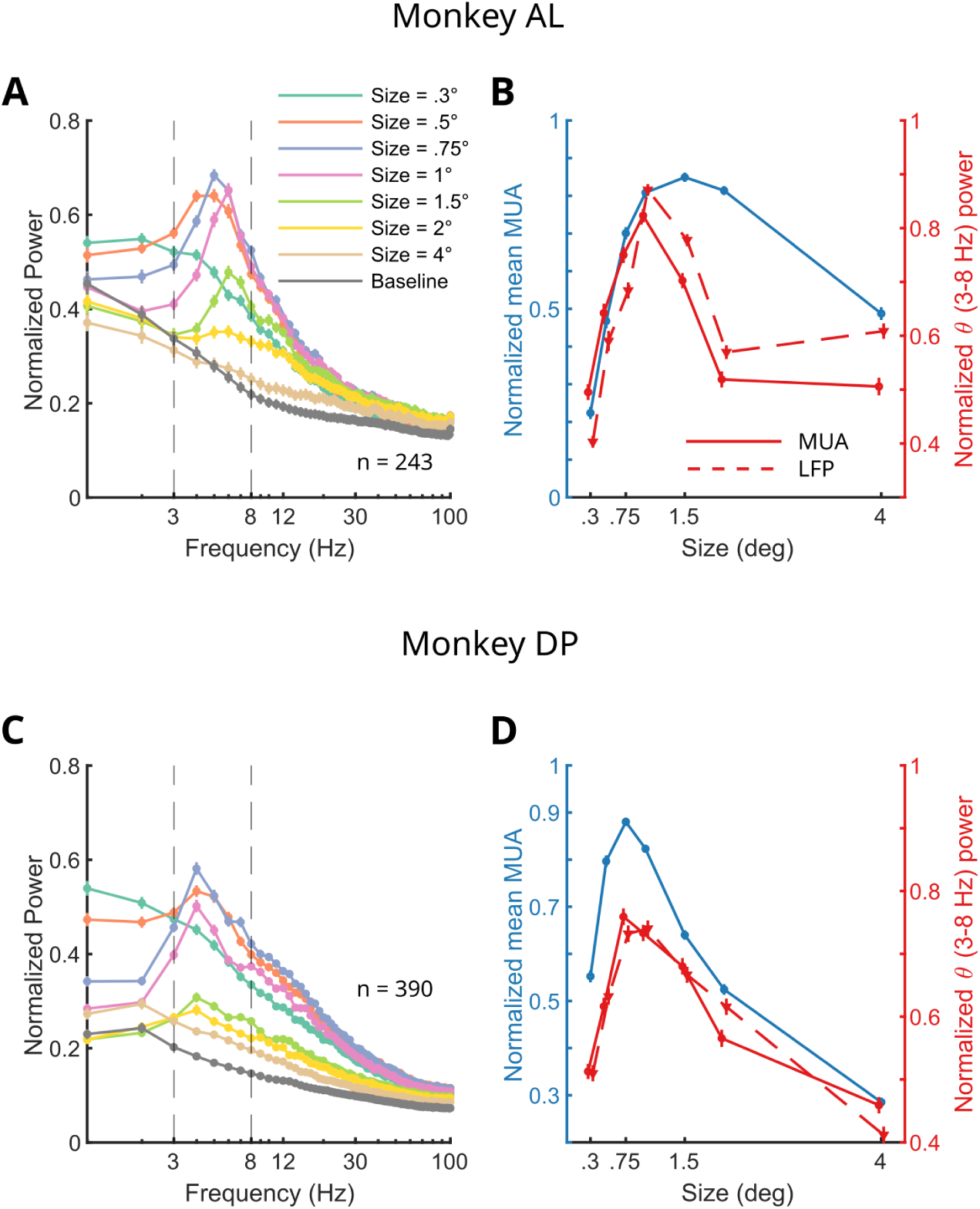
Theta oscillations at different stimulus sizes: population results. (A) Normalised population power spectra of MUA for each size averaged across channels for monkey AL (n = 248 channels). Dashed lines delineate theta frequency range. Error bars are ± 1 standard error of the mean (SEM). (B) Normalised mean MUA (blue), MUA theta power (red continuous line), and LFP theta power (red dashed line) across different sizes averaged across all channels for monkey AL. Error bars are ± 1 SEM. (C) Same as A, but for monkey DP (n = 390 channels). (D) Same as B, but for monkey DP.

Having established the dependence of theta oscillation on stimulus size, we tested the impact of a second stimulus property known to modulate V1 response, i.e. contrast. Responding to different contrast intensities is another fundamental function of the visual system (Albrecht & Hamilton, 1982). To test this, we stimulated the receptive field with a black disk with varying contrast levels relative to its background. The size of the black disk was optimized to elicit theta oscillations from a size-tuning test conducted beforehand. Similar to size tuning, higher stimulus contrast increased MUA and theta power (Supplementary Figures 9 and 10).

After confirming the stimulus dependence of V1 theta oscillations, we explored whether this rhythmic activity is related to behavior. Given prior associations between neural theta oscillations and microsaccades—miniature eye movements occurring approximately every 250 ms that signal active exploration and cognitive engagement (Bosman et al., 2009; Lowet et al., 2016; Otero-Millan et al., 2008)—we investigated whether microsaccades could account for the observed theta oscillations. Using a standard detection algorithm (Engbert & Kliegl, 2003), we analyzed microsaccades during the size-tuning task (Supplementary Figure 4). However, our analyses did not support a convincing relationship between microsaccades and theta oscillations, suggesting that microsaccades do not explain the emergence of theta oscillations in V1 under our experimental conditions (Supplementary Figures 5–8).

To further explore the behavioral relevance of stimulus-induced V1 theta oscillations, we trained monkeys to perform a visual detection task (Figure 3A). Monkeys fixated on a central spot while a visual stimulus, known to induce theta oscillations in V1 MUA, was presented. Monkeys were required to detect a subtle luminance change at the stimulus center and report it by making a saccade toward the change location. Critically, the timing of the luminance change varied between 500 and 1500 ms after stimulus onset, allowing assessment of reaction times (RTs) at different phases of the neural theta rhythm. Previous research in humans and monkeys has established theta-rhythmic performance measures in similar detection tasks (Fiebelkorn et al., 2013, 2018; Kienitz et al., 2018; Landau & Fries, 2012).

Plotting RTs against visual target onset times from one session revealed a pattern of alternating shorter and longer RTs (Figure 3B). Spectral analysis indicated that RTs fluctuated at a theta frequency of 6 Hz (Figure 3C, blue). Correspondingly, MUA oscillated at a theta frequency peaking at 5 Hz. Across sessions in both monkeys, there was a moderate positive correlation between the peaks of MUA and RT power spectra (r = 0.547 [95% CI 0.118–1], p = 0.021, one-tailed, n = 14; Pearson correlation), suggesting that stimulus-induced theta rhythms in V1 are reflected in behavioral performance fluctuations.

Previous studies have reported correlations between behavioral performance and the phase of theta oscillations in higher-level brain areas of humans and macaques (Busch et al., 2009; Dugué et al., 2015; Fiebelkorn et al., 2018; Helfrich et al., 2018; Kienitz et al., 2018). To assess whether the phase of V1 theta oscillations correlates with RT, we calculated theta phase in hit trials of the detection task (Figure 3). We sorted trials by RT, and intertrial phase coherence (ITPC) (Tallon-Baudry et al., 1996) was computed for fast, slow, and combined (fast+slow) trials. Fast and slow trials exhibited phase clustering in opposite directions, while combined trials did not, indicating that different RTs were associated with specific theta phases (Figure 4A). The phase opposition sum (POS) (VanRullen, 2016) was calculated for each channel and time point around target presentation (-300 ms to 150 ms) (Figure 4B, top). Statistical analysis using bootstrapping revealed significant POS values approximately 200–250 ms before target presentation, suggesting that the phase of V1 theta oscillations influences RTs (Figure 4B, bottom).

## Discussion

Our study reveals that specific visual stimuli can induce theta-band (3–8 Hz) oscillations in the spiking activity and local field potential (LFP) of primary visual cortex (V1) of macaque monkeys, correlating with visual detection. This aligns with prior research in rodents and primates, highlighting neural theta rhythms in V1 during sensory processing and active exploration (Fournier et al., 2020; Gao et al., 2021; Kienitz et al., 2021; Niell & Stryker, 2010; Spyropoulos et al., 2018; Zimmerman et al., 2025; Zold & Hussain Shuler, 2015).

Our findings extend these observations by demonstrating that the emergence of theta oscillations in V1 depends on basic visual stimulus properties. Consistent with studies in the primate visual association cortex (Kienitz et al., 2018; Rollenhagen & Olson, 2005), we emphasize the role of receptive field (RF) computations in generating theta rhythmic activity. We observed that theta oscillations correlate with spike rates; stimuli that effectively drove neural populations also induced stronger theta oscillations. This suggests that in V1, theta oscillations are inherent to the spiking patterns of neural populations. Minimal activation, such as during non-optimal stimuli or pre-stimulus periods, results in weaker theta oscillations, whereas optimal stimuli enhance them.

Many V1 neurons exhibit a preference for stimuli drifting at theta frequencies (Allison et al., 2001; Foster et al., 1985; Hawken et al., 1996; Yu et al., 2010), indicating that these neurons may function as temporal filters tuned to theta frequencies. Thus, sustained presentation of a static stimulus, as in our study, could elicit rhythmic firing at theta frequency, as it reflects the inherent filter properties of these neurons. Similarly, locomotion in rodent studies might effectively drive neural activity aligned with their preferred theta filtering. Stronger stimulus activation correlates with increased theta activity, aligning with our observations across different stimulus properties.

The presence and modulation of stimulus-induced theta oscillations in V1 by specific stimulus properties suggest that theta rhythms are crucial in visual processing. They may provide a temporal framework for organizing sensory information, thereby enhancing visual perception efficiency. This supports the notion that perceptual exploration operates at a theta rhythm, as indicated by psychophysical research in humans (Dugué et al., 2015, 2016; Fiebelkorn et al., 2013; Kienitz et al., 2022; Landau & Fries, 2012; Michel et al., 2021; Re et al., 2019; Song et al., 2014). Moreover, the correlation between theta phase in V1 and behavioral performance suggests that rhythmic exploration may originate from visual cortical processing. This complements earlier studies indicating that rhythmic activity and behavior result from large-scale interactions between higher-level brain areas (Dugué et al., 2019; Fiebelkorn et al., 2018; Fiebelkorn & Kastner, 2021; Gaillard et al., 2020; Helfrich et al., 2018).

In summary, our study demonstrates that specific visual stimuli elicit theta-band oscillations in macaque V1, which are functionally relevant to visual detection performance. These findings bridge observations from rodent models to human research, emphasizing the conserved role of theta rhythms in visual processing and their significance in shaping perceptual experiences.

### Theta oscillations are related to behavioural performance

**Figure 3.**
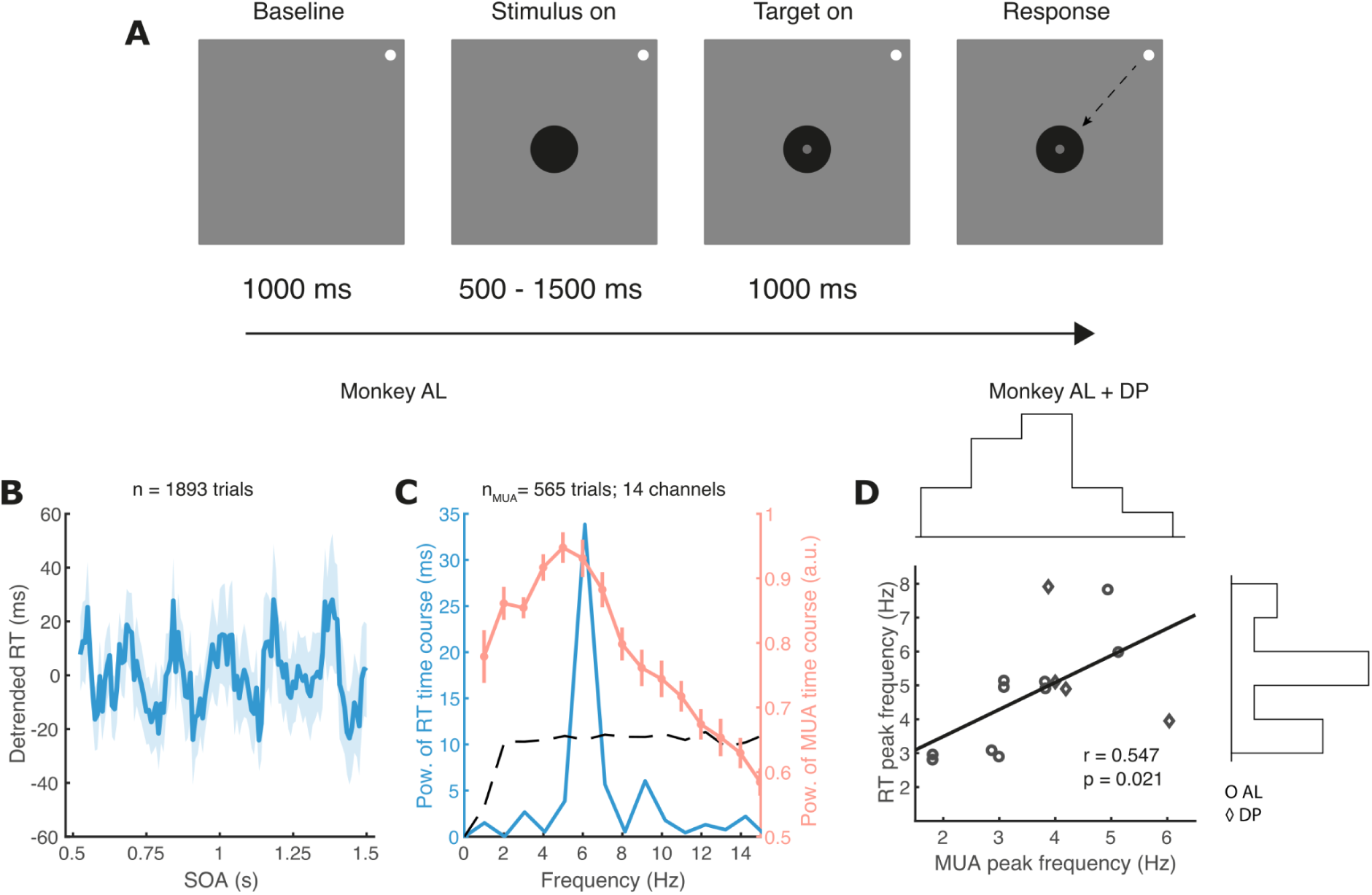
Theta oscillations at different stimulus sizes: population results. (A) Behavioural task. The monkeys had to detect a small luminance change (grey dot), in the centre of a stimulus (black disk) and make a saccade toward the stimulus. The interval between stimulus and luminance change, stimulus onset asynchrony (SOA), was varied densely between 500 to 1500 ms. (B) Reaction times (RT) as a function of SOA from one session. (C) Power spectrum of the time course of the RT in B (blue), superimposed with the average neural power spectra recorded during the task (orange). Black dashed line is the threshold of statistical significance for the RT power spectrum after correction for multiple comparisons (α = 0.05 / 15 frequencies = 0.0033). (D) Scatter plot showing the relationship between peak frequency of MUA power spectra and peak frequency of RT power spectra (n = 14 sessions). Sessions are combined for both monkeys. We introduced a small jittering to the data points in the scatter plot for illustration purposes only to improve visibility because of the overlaps of some data points. The correlation coefficient and regression line were calculated from the non-jittered data. There was a significant positive relationship between MUA peak frequency and RT peak frequency. Marginal histograms showed the concentration of peak frequency of both MUA (top histogram) and RT (bottom histogram) in the theta range.

**Figure 4.**
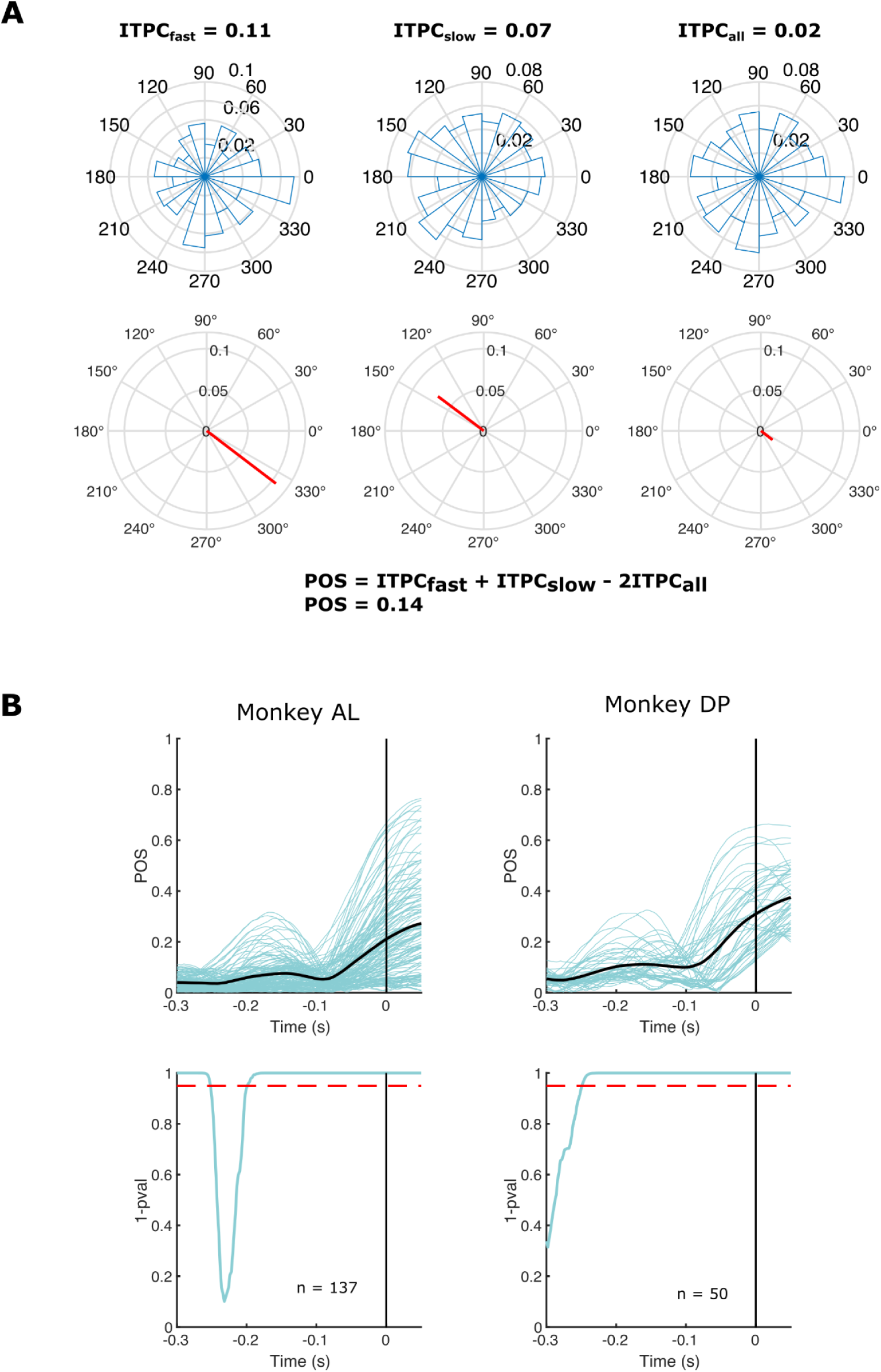
The phase of theta oscillations is correlated with reaction times. (A) An illustration of Phase Opposition Sum (POS) calculation from an example channel at one time point. We calculated Intertrial Phase Coherence (ITPC) for fast, slow, and fast + slow trials. The top row shows the phase distribution across trials, while the bottom row shows the ITPC (red lines). The line direction shows the mean phase angle, while the length shows how tightly the phase distribution clusters. POS was calculated according to the equation at the bottom. A larger value means a larger ITPC difference between fast and slow trials, implying a stronger correlation between theta phase and RT. (B) Population POS across channels (top) and p-value (bottom). We calculated POS for every channel and every time point around the target presentation (t = 0s, vertical line). Blue lines are the POS of individual channels, and the black line is the mean POS across channels. P-value was calculated for every channel and combined (bottom row). Note that we plotted the p-value as 1 - p-value. The red dashed line is the significance threshold (p = 0.05, corrected for false discovery rate (FDR).

## Methods

### Subjects

Two healthy adult female rhesus monkeys (*Macaca mulatta*, monkey AL: age 5 years old and weight 8 kg; monkey DP: age 6 years old and weight 9 kg) participated in this study. We implanted a head post and a recording chamber over area V1 in the right hemisphere. Anaesthesia procedure, surgical procedure, implant methods, and postoperative conditions are described in a previous publication (Ortiz-Rios et al., 2018). During the testing period, the monkeys were put under a fluid control procedure which did not impair the animals’ physiology and welfare (Gray et al., 2016). All procedures complied with UK Animals Scientific Procedures Act 1986 and European Council Directive 2010/63/EU.

### Neurophysiological recordings

Neurophysiological data were collected by two types of electrodes used in different experimental sessions: (1) single tungsten electrodes with epoxylite coating (FHC, Bowdoin, USA) and (2) silicone linear probes with 16 (1 shaft) or 32 channels (2 shafts with 200 µm spacing between the shafts), 150 µm inter-electrode spacing, with platinum contacts (Atlas Neuroengineering, Leuven, Belgium). Single FHC electrodes were referenced to the stainless-steel guide tube used to penetrate the dura, while Atlas linear probes were referenced to a silver wire placed on the dura while the chamber was filled with saline. Electrodes were inserted daily into the right V1 of each monkey with a hydraulic micromanipulator (Narishige, Japan). Raw data were recorded at a sampling rate of 30 kHz using a Blackrock Microsystems Cerebus system (Blackrock Microsystems, Utah, USA).

Neurophysiological data were analysed using MATLAB-based NPMK software (Blackrock), custom-written MATLAB codes (Mathworks), and the FieldTrip MATLAB software toolbox (Oostenveld et al., 2011). The envelope of multi-unit activity (MUA) (Drebitz et al., 2019; Supèr & Roelfsema, 2005) was obtained by high-pass filtering (8th-order Chebyshev filter with 300 Hz cutoff frequency) the raw data, rectifying (taking the absolute values), and downsampling to 500 Hz. LFP signal was obtained by down-pass filtering the data (7th-order Chebyshev filter with 150 Hz cutoff frequency), followed by downsampling to 500 Hz. To remove common noise in the LFP, we re-referenced the data with the common average referencing method. This was performed by subtracting the average activity across all channels from every channel before the data was cut into trials. The re-referencing was only performed on data collected during passive fixation, not on data collected during behavioural task.

### Visual stimulation

Stimulus presentation and monkey behaviour were controlled by MWorks (https://mworks.github.io). During the experiments, eye movements were tracked monocularly and recorded using an infrared-based eye-tracking system with 500 Hz sampling rate (EyeLink 1000, SR Research, Ottawa, Canada). Stimuli were presented with a ViewPixx LCD monitor with a 120 Hz refresh rate, 1920 x 1080 pixels resolution, and 24-inch diagonal display size (VPixx technologies, Saint-Bruno, Canada). The monitor luminance output was linearized by measuring the luminance of red, green, and blue using a photometer at 8 brightness levels repeated 10 times each. We fitted the luminance profile with a power function and applied the inverse of the power function during stimulus presentation. The viewing distance was 85 cm.

RF location was estimated by presenting a black square (luminance 0.1 cd/m2) in quick succession, 100 ms each, at various spots on the screen with a grey background (luminance 45 cd/m2). The animals fixated a white fixation dot (0.3° diameter, 92 cd/m2 luminance). Usually, we started by presenting a 1° wide square in a non-overlapping 5 by 5 grid to find the rough location of the RF. The MUA responses to the RF stimuli were analyzed to find the spot where activation was highest. We considered this spot to be the centre of the RF location. Once we confirmed the rough RF location, the procedure was repeated with 0.5° and 0.25° squares to further delineate the RF location at a higher spatial resolution. The RF centre location obtained with 0.25° stimulation was recorded and subsequent visual stimulation was always placed at this location. In the case where we recorded with multiple electrodes, we usually focused on the RF location of one or two channels with the strongest activation as judged by listening to the auditory-converted neural activity and/or by visual inspection of the MUA data from RF analysis.

For the passive viewing task, the animals were required to maintain fixation on a small white dot while various stimuli were presented at the receptive field (RF). All stimuli were always presented with a gray background (RGB [0.5,0.5,0.5] luminance 45 cd/m2). Trials were initiated by the animals looking at the fixation dot followed by a fixation time of at least 1000 ms. After that, the stimuli were presented for 1200 ms. Animals were rewarded if they maintained fixation for the whole duration of the trial. If the animals moved their eyes outside the fixation window any time before the stimulus disappeared, the trials were aborted, and they weren’t rewarded. The stimulation conditions from failed trials were included again randomly in the protocol to ensure adequate testing across the different task conditions. The radius of the fixation window was typically 1°. Depending on the animals’ condition and motivation, the radius could be smaller, but never exceeded 1°.

To test the effect of stimulus size on theta oscillations, we presented a black (RGB [0,0,0] luminance 0.1 cd/m2) disk with one of 7 possible sizes (0.3°, 0.5°, 0.75°, 1°, 1.5°, 2°, and 4° diameter). The size of the stimulus varied on a trial-by-trial basis. For most of the recordings, we presented each size for 40 trials; however, in some sessions, we presented more trials per size depending on the animals’ motivation. After the size tuning recording was finished, the data was analyzed to identify the size which induced the strongest theta power. This size was recorded and used to present stimuli for the contrast and orientation experiment.

To test the effect of contrast, we presented a grey disk with varying contrast relative to the grey background. On a single trial, we presented the disk with 8 possible contrasts (5.3 %, 9.9 %, 14.6 %, 20.4 %, 25.4 %, 30.4 %, 49.5 %, 100). The contrast was calculated as Michelson contrast and varied every trial. For most recordings, we presented each contrast for at least 40 trials. For one session in monkey DP, we presented only 20 trials for each contrast. In some sessions, depending on the motivation of the monkey, we presented more than 40 trials per contrast.

For the detection task, the monkeys maintained fixation for 1000 ms before a stimulus was presented in the RF. The stimulus was a black disk (luminance 0.1 cd/m2) on a grey background (luminance 45 cd/m2). The stimulus size was optimized to generate theta oscillations from a size tuning passive viewing task performed before the detection task. The stimulus size that induced the strongest theta oscillations was used as the stimulus size in the detection task. After a random 500 to 1500 ms of stimulus presentation, a target appeared in the centre of the stimulus and stayed on the screen for a maximum of 1000 ms. The monkeys had to detect the target by making a saccade toward the stimulus within 1000 ms after target onset. A successful saccade to the stimulus was rewarded. Fixations break from the fixation point at any time before target onset would abort the trial. In 25 % of trials, we presented catch trials where there was no target presentation. In the catch trials, the monkeys were rewarded for maintaining fixation for 1500 ms.

### Data analysis

#### Passive viewing task

Our main unit of analysis is the MUA collected per electrode for the FHC electrodes and MUA or LFP per electrode channel for the linear probes. For our main analysis, we only included visually responsive channels. To determine the visually responsive channels, we compared the 100 ms post-stimulus onset MUA to the last 500 ms stimulus baseline before stimulus onset. We did this calculation separately for each stimulus domain, size and contrast. Visually responsive channels were the channels with activity higher than 2.5 standard deviations of mean baseline activity in at least 4 stimulus conditions (e.g., 4 sizes or 4 contrasts). The channel selection for LFP analysis was based on the MUA channel selection.

Frequency analysis was performed using the ft_freqanalysis function from FieldTrip toolbox. The fast fourier transform (FFT) analysis was performed at every channel and at the single trial level with single taper Hanning window (cfg.method = ‘mtmfft’ and cfg.taper = ‘hann’ as an input for ft_freqanalysis). We extracted the power of frequency 1 - 100 Hz from the time period 200 ms - 1200 ms post-stimulus. Baseline power spectra were calculated with the same method from period -1000 ms to 0 ms pre-stimulus. The power spectrum of each channel was obtained by averaging the single-trial power spectra for the tested stimulus condition.

For analysis at the population level, the power spectra from every channel were peak normalized with the following method: first, we identified the strongest power value from all trial-averaged spectra across stimulus conditions. Second, we divided all power spectra to the power from the overall strongest spectrum. This peak normalization procedure also ensured that our results were not disproportionately affected by some channels with particularly strong power values.

For the peak frequency analysis, we calculated the mean power spectra across trials for every channel and identified the peak with the strongest power. The distribution of peak frequency at all channels was plotted using a MATLAB function plotSpread (Jonas, 2025)(Jonas, 2021).

To construct tuning curves, we calculated the mean MUA from the same time window we used for frequency analysis, 200 ms - 1200 ms post-stimulus. Similar to the frequency analysis, we also performed peak normalization of tuning curves for the population level analysis. For every channel, we identified the stimulus condition with the highest MUA and divided the MUA values at other conditions by the maximum value.

#### Microsaccades analysis

We detected microsaccades using an algorithm developed by Engbert and Kliegl (Engbert & Kliegl, 2003). The raw eye traces were first converted to velocities with a moving average window of 5 data samples. Then, on each trial, we computed the threshold of microsaccades detection as 6 times the standard deviation of velocity time series. An event was classified as microsaccades if it exceeded the threshold by at least 3 data samples (minimum duration of 6 ms). We removed all microsaccades with amplitude larger than 1°.

#### Detection task

The analysis of behavioural data began with removing trials with RT less than 100 ms or larger than 1000 ms. Next, we binned the trials based on the SOA between stimulus and target. The bin was a 50 ms sliding window with an 8.3 ms step. Mean RT was calculated as a function of binned SOA to generate the RT time course. Then we detrended the time course by fitting a second-order polynomial function and removing the fit from the RT time course. We then calculated the FFT of the detrended time course to see if the resulting power spectrum contains a peak in the theta range. To establish statistical significance, we generated a surrogate time course by randomising the RT and SOA from the real, non-detrended time course. This randomisation destroyed the temporal structure, from the original time course. After that, we detrended and calculated FFT from the surrogate time course. This analysis was repeated for 5000 times and for each iteration we obtained the power spectra from the surrogate time course. This results in a surrogate distribution of power value from each frequency. We then set the threshold for statistical significance by taking the 99.67 percentile value from the surrogate distribution for each frequency. The percentile value was corrected for multiple comparisons across 15 frequencies (1-15 Hz), 1 - (0.05/15) = 99.67.

Similar to what we did for the passive viewing task, we analysed the MUA only from visually responsive channels, defined as channels with the first 100 ms MUA higher than at least 2.5 SD of the last 500 ms of baseline. The frequency analysis for the detection task was performed on 500 - 1500 ms after stimulus onset on successful catch trials only, which were trials where the target did not occur and the animals maintained fixation for 1500 ms. The reason why we only analyzed the catch trials was to obtain the full stimulus-induced MUA, uninterrupted by the response to target onset and the saccadic response to it.

We tested the relationship between behavioural and neural oscillations by analyzing both measures across sessions (n = 14), combined for both monkeys (n = 10 for monkey AL and n = 4 for monkey DP). For each session, we extracted the peak frequency, the frequency with the highest power, separately from RT and MUA power spectrum. All RT peak frequencies in each session were statistically significant as defined above. The peak frequency of MUA was obtained from the mean peak normalized power spectra across channels. We then calculated Pearson correlation coefficient between the RT and MUA peak frequency.

To test the correlation between theta oscillations and behavioural performance, we used the LFP signal on hit trials where the monkey correctly made a saccade toward the stimulus after target onset. We bandpass-filtered the LFP in the theta range, 3 - 8 Hz, with a windowed-sinc FIR filter and performed a Hilbert transform to extract the instantaneous phase (cfg.filttype = ‘firws’ and cfg.hilbert = ‘angle’ in ft_preprocessing function of FieldTrip toolbox). Next, we sorted our behavioural trials based on RT. We obtained the fastest and slowest 25 % of the trials and classified them as fast and slow, respectively.

After that, we calculated the Phase Opposition Sum (POS) values following the steps laid out by VanRullen (VanRullen, 2016) (see Figure 4 for example): First, we calculated the ITPC for the fast, slow, and fast+slow trials. Second, POS was defined as POS = ITPC_fast_ + ITPC_slow_ - 2ITPC_all._ POS was calculated for every time point from -300 ms before and 50 ms after the target presentation and for every channel. Third, we performed a non-parametric permutation procedure to test for statistical significance. In the permutation test, assigning phase and RT from random trials destroyed the relationship between theta phase and RT. This was done 1000 times, and the POS values were calculated for each permutation to obtain the null POS distribution. Next, we compared the experimental POS values with the POS null distribution and expressed it as a z-score. The z-score value was converted to a value using the inverse cumulative normal distribution function. The permutation test was performed at every time point and channel. Last, the list of p-values from all channels was combined using the Fisher method (Fisher, 1938; Yoon et al., 2021) and corrected for multiple comparisons (p < 0.05) using the False Discovery Rate. We calculated the POS and converted it to p-values using MATLAB scripts provided by VanRullen (VanRullen, 2016).

## Acknowledgement

This work was supported by starting grant OptoVison 637638 and SNF grants BSET-0_201532, 31NE30_203974 and 310030_204544 to Michael C Schmid. We thank the Comparative Biology Centre for their help with animal handling and training, Michael Ortiz-Rios for help with surgeries, and Ian Milne for technical assistance for this project. Alex Thiele, Chris Petkov, Alex Maier, Alwin Gieselmann, Samy Rima, and J. Anthony Movshon for his constructive comments.

## Supplementary Figures

**Supplementary Figure 1.**
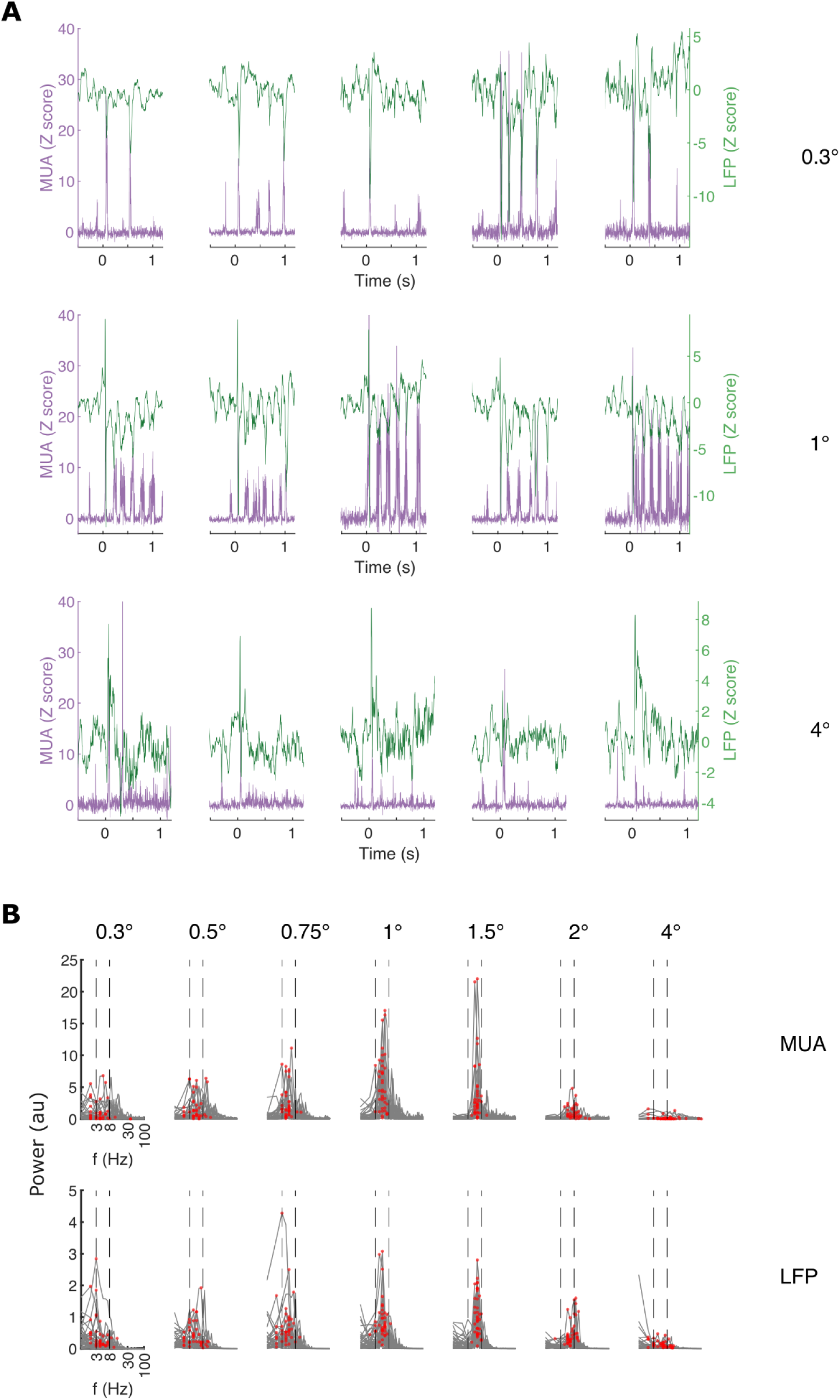
Example single trial time course and power spectra. (A) Five example trials of MUA (purple, left y-axis) and LFP (green, right y-axis) response to three different stimulus sizes on each row. The signals were obtained from the same channel in Figure 1. Each column corresponds to one trial. Oscillatory activity is clearly seen for 1° stimulus size for both MUA and LFP. (B) Single-trial power spectra (grey) of post-stimulus MUA (top) and LFP (bottom). Dashed lines delineate the theta frequency band. The red dots are peak frequencies for every trial.

**Supplementary Figure 2.**
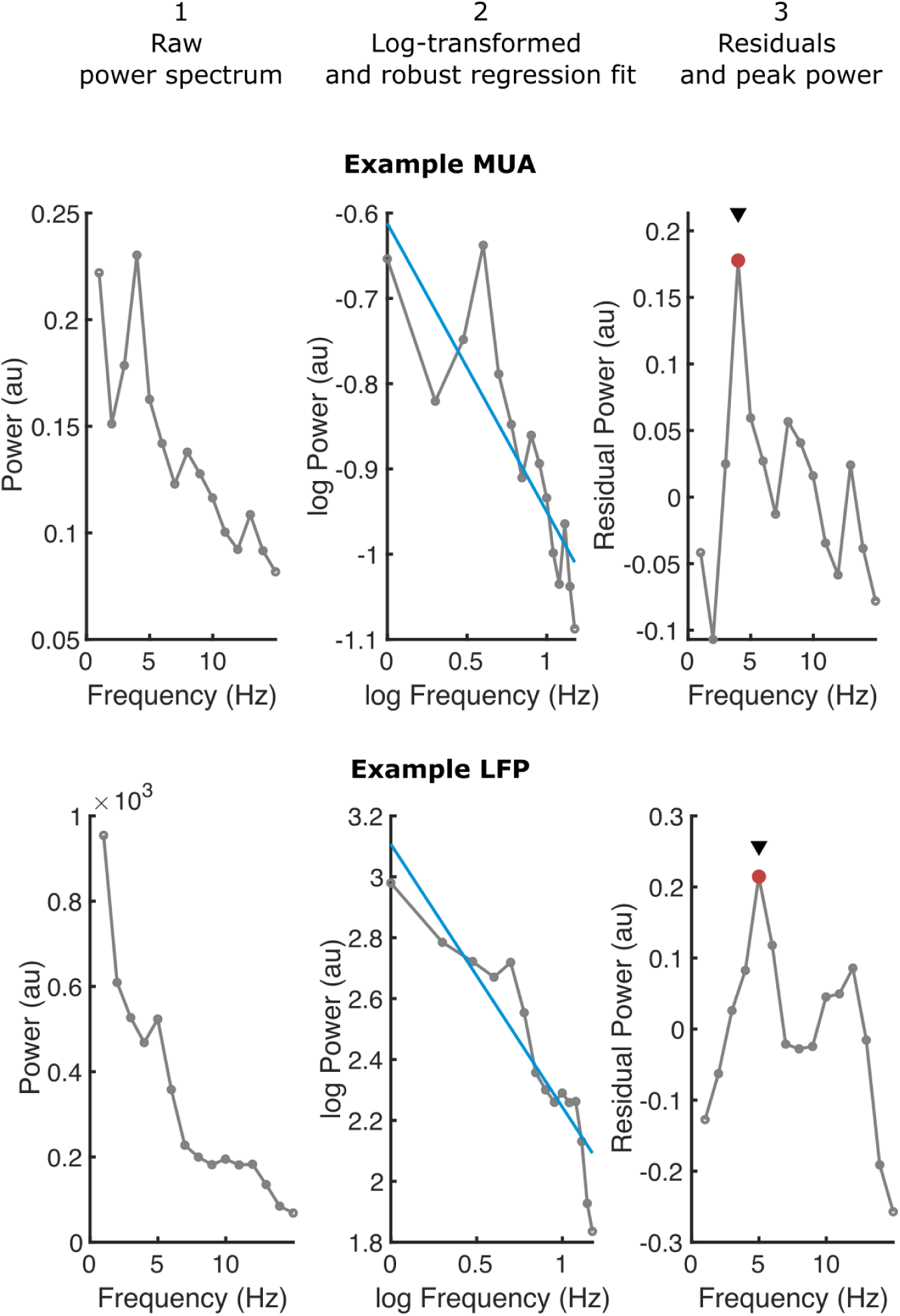
Method to calculate theta power. To calculate theta power, we first limited the power spectra in the 1 - 15 Hz range (step 1). Next, we log-transformed both the x (frequency) and y-axis (power) and fit a robust regression model (step 2). Last, we removed the linear trend and obtained the residual power (step 3). Theta power was the largest peak in the residual spectra.

**Supplementary Figure 3.**
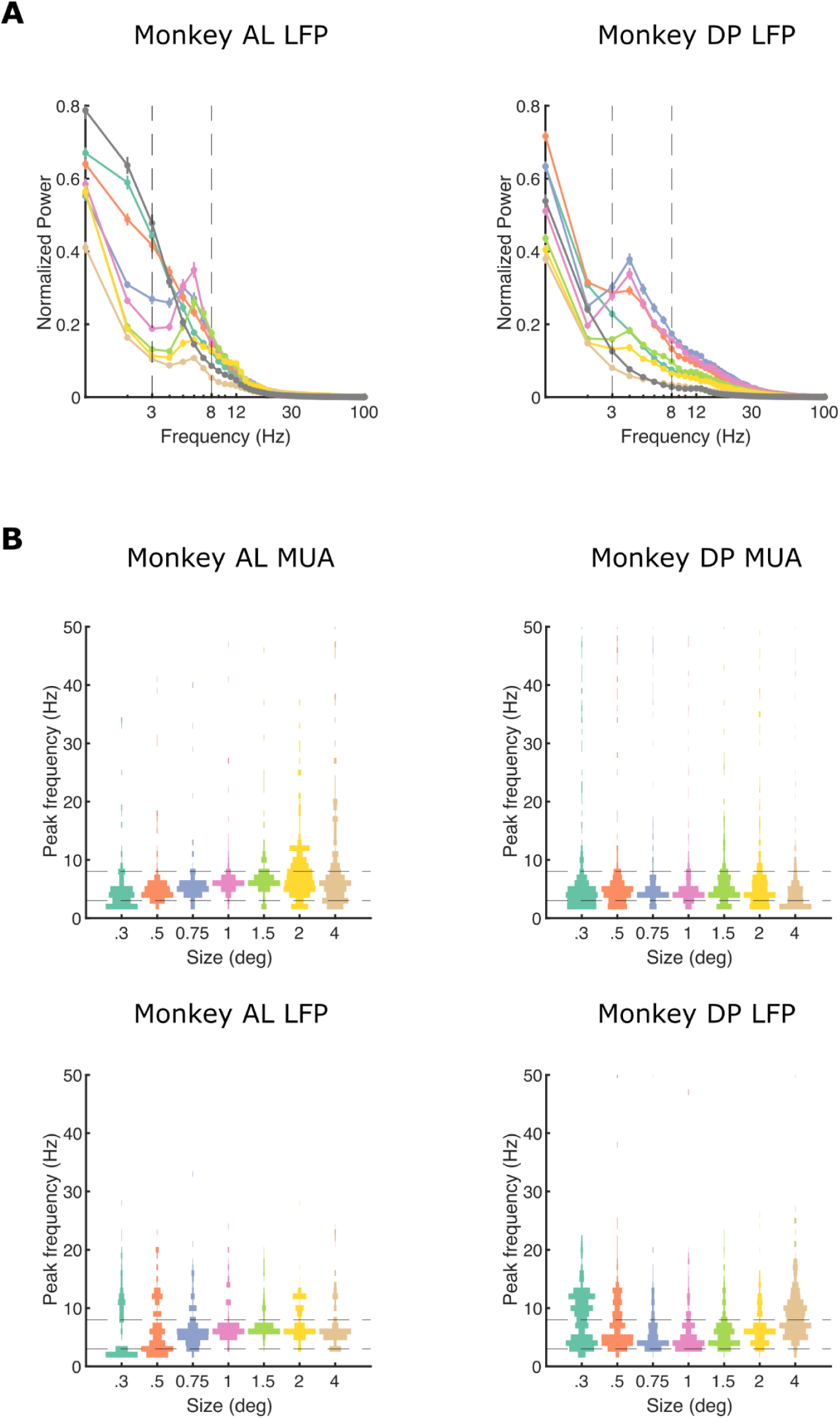
(A) Normalised population LFP power spectra for each size averaged across channels for monkey AL (n = 248 channels) and DP (n = 390 channels). Dashed lines delineate theta frequency range. Error bars are ± 1 standard error of the mean (SEM). (B) Peak frequency distribution across channels at different stimulus sizes for each monkey, for MUA (top) and LFP (bottom) separately. We calculated the mean power spectra across trials for every channel and identified the peak with the strongest power. Theta frequency band is delineated by horizontal dashed lines.

**Supplementary Figure 4.**
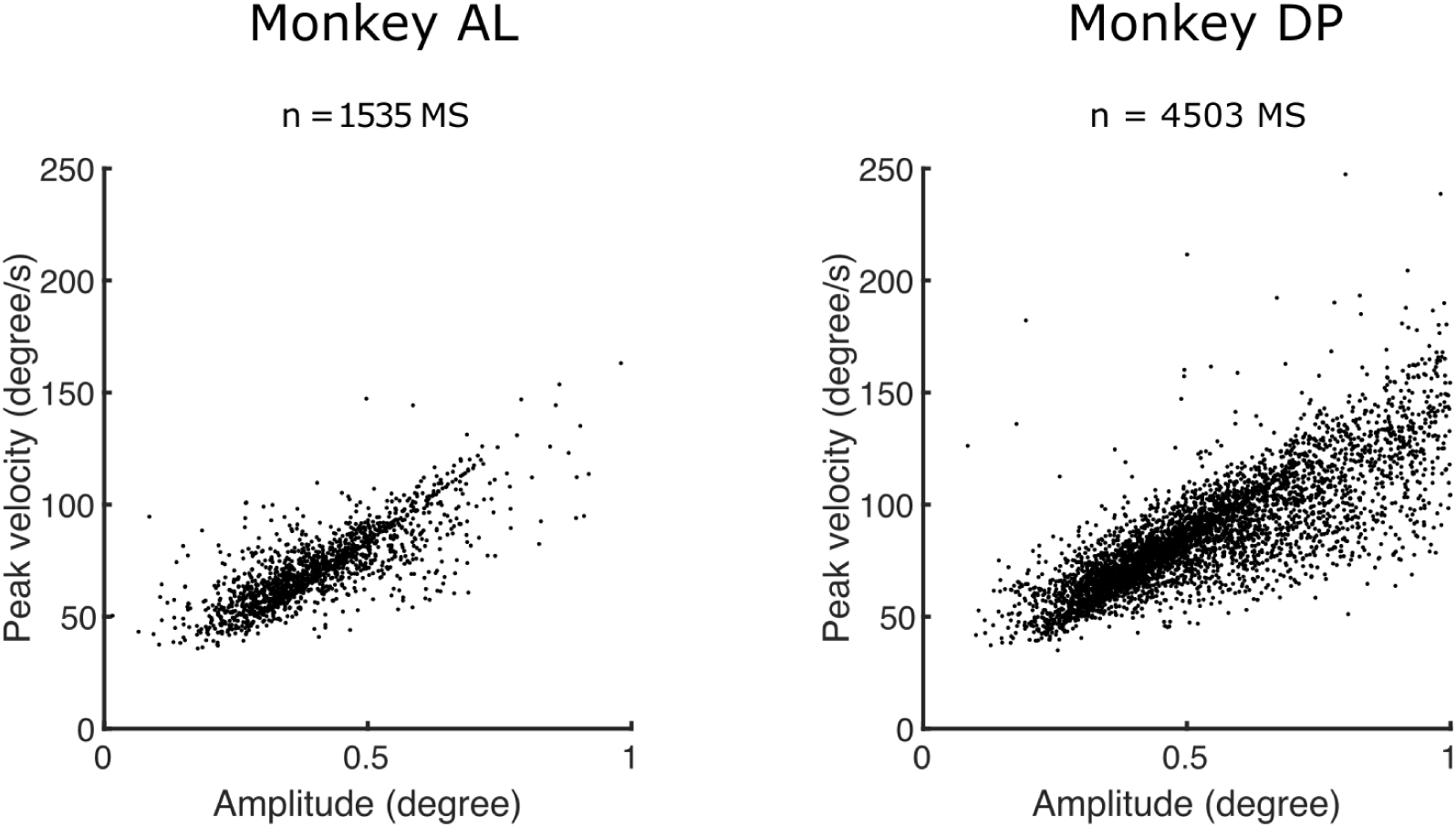
Microsaccades main sequence showing the positive relationship between amplitude and peak velocity.

**Supplementary Figure 5.**
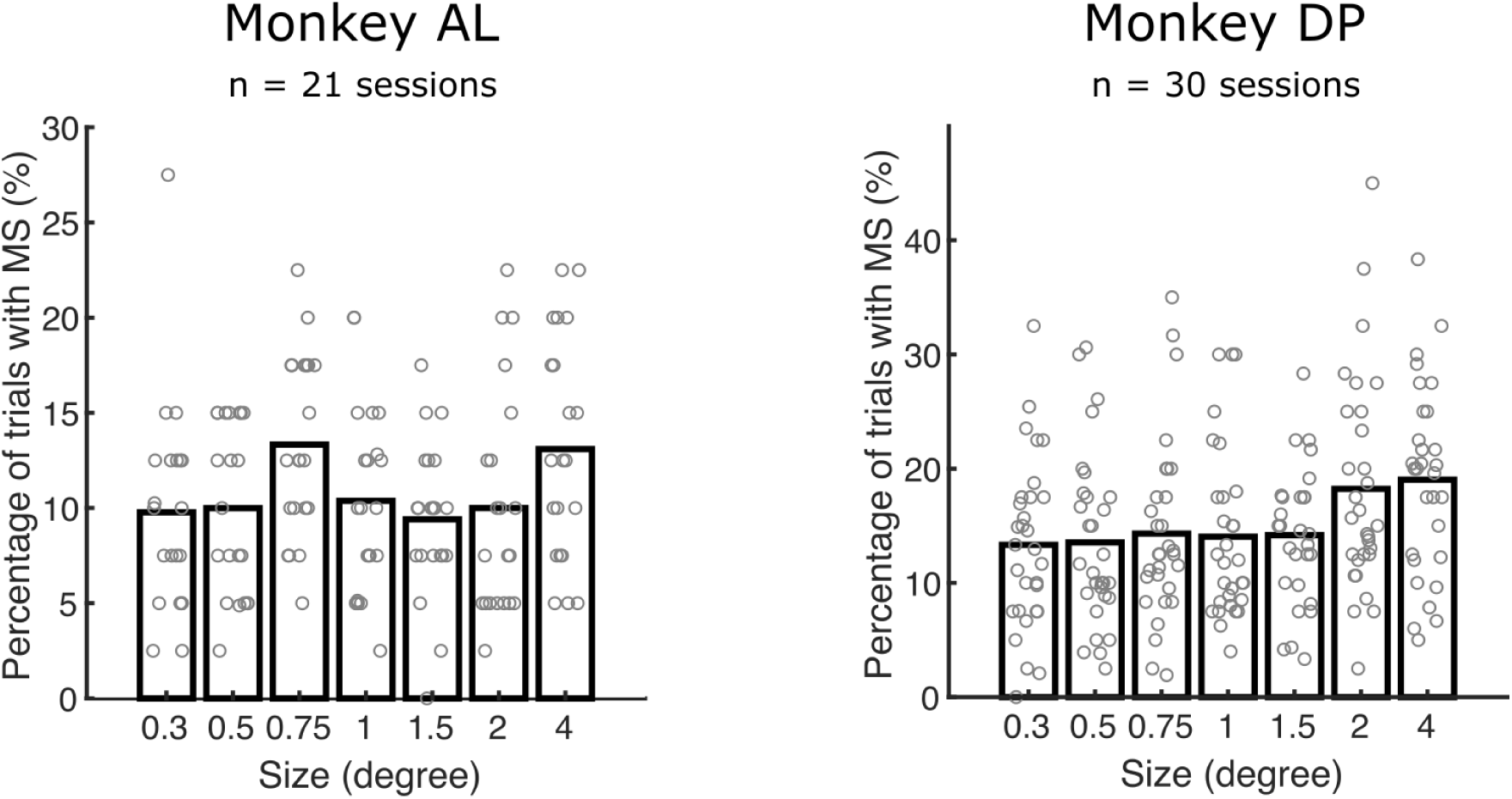
Mean percentage of trials with microsaccades across stimulus sizes. Each dot represents a single session. Microsaccades were rare events. Microsaccades occurred on ∼ 10 - 15 % of trials across sizes. This microsaccades occurrence pattern differed substantially from our observations that post-stimulus neural theta oscillations are consistently present across trials (Supplementary Figure 1 as an example).

**Supplementary Figure 6.**
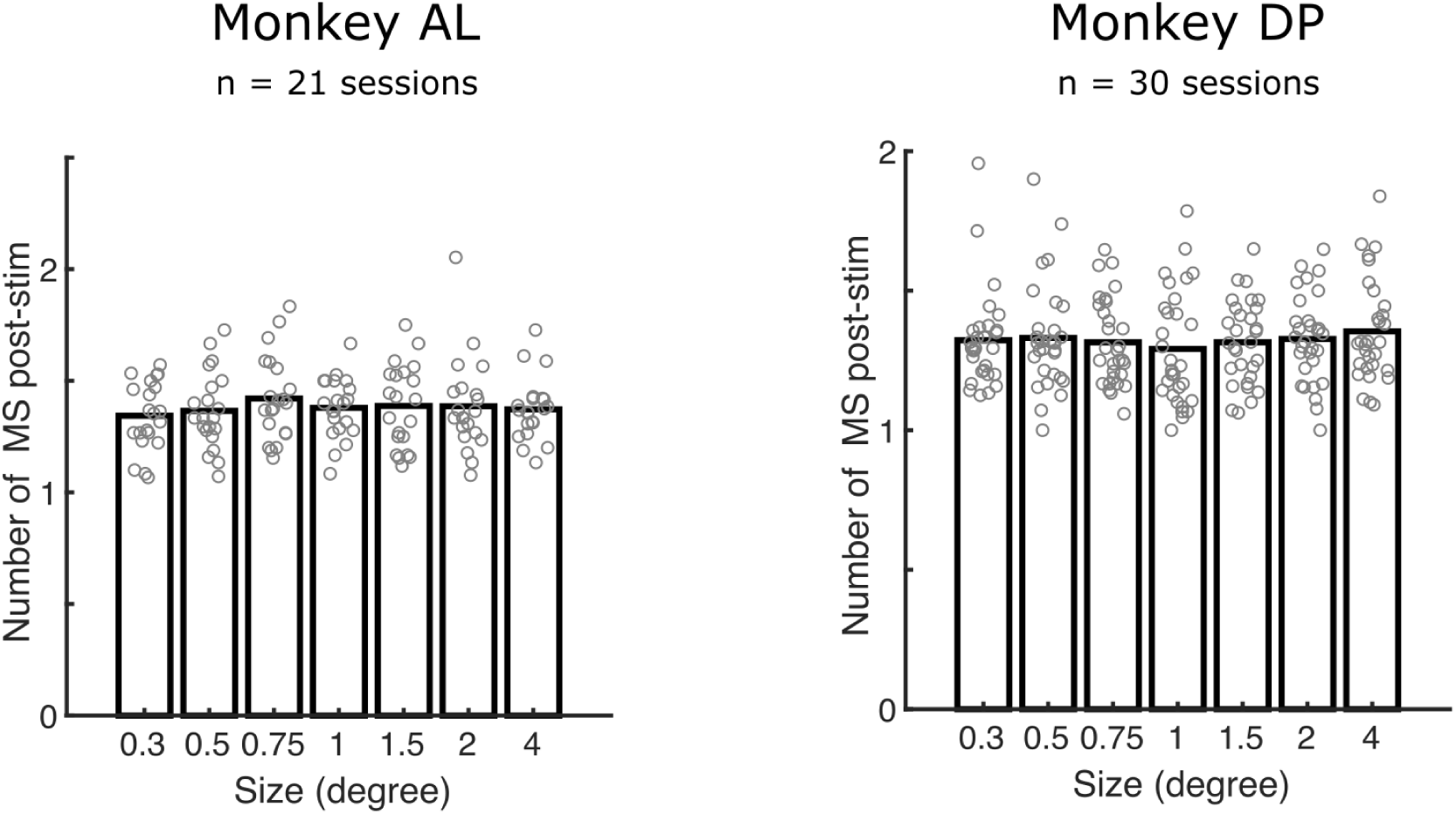
Number of microsaccades in the post-stimulus period in trials where microsaccades occurred after stimulus onset. Each dot represents a single session. On average, microsaccades occurred once during the whole post-stimulus period. The number of microsaccades during the post-stimulus period was not different across stimulus sizes (p > 0.05, Friedman test). These findings are inconsistent with rhythmic saccades inducing theta rhythmic neural activity.

**Supplementary Figure 7.**
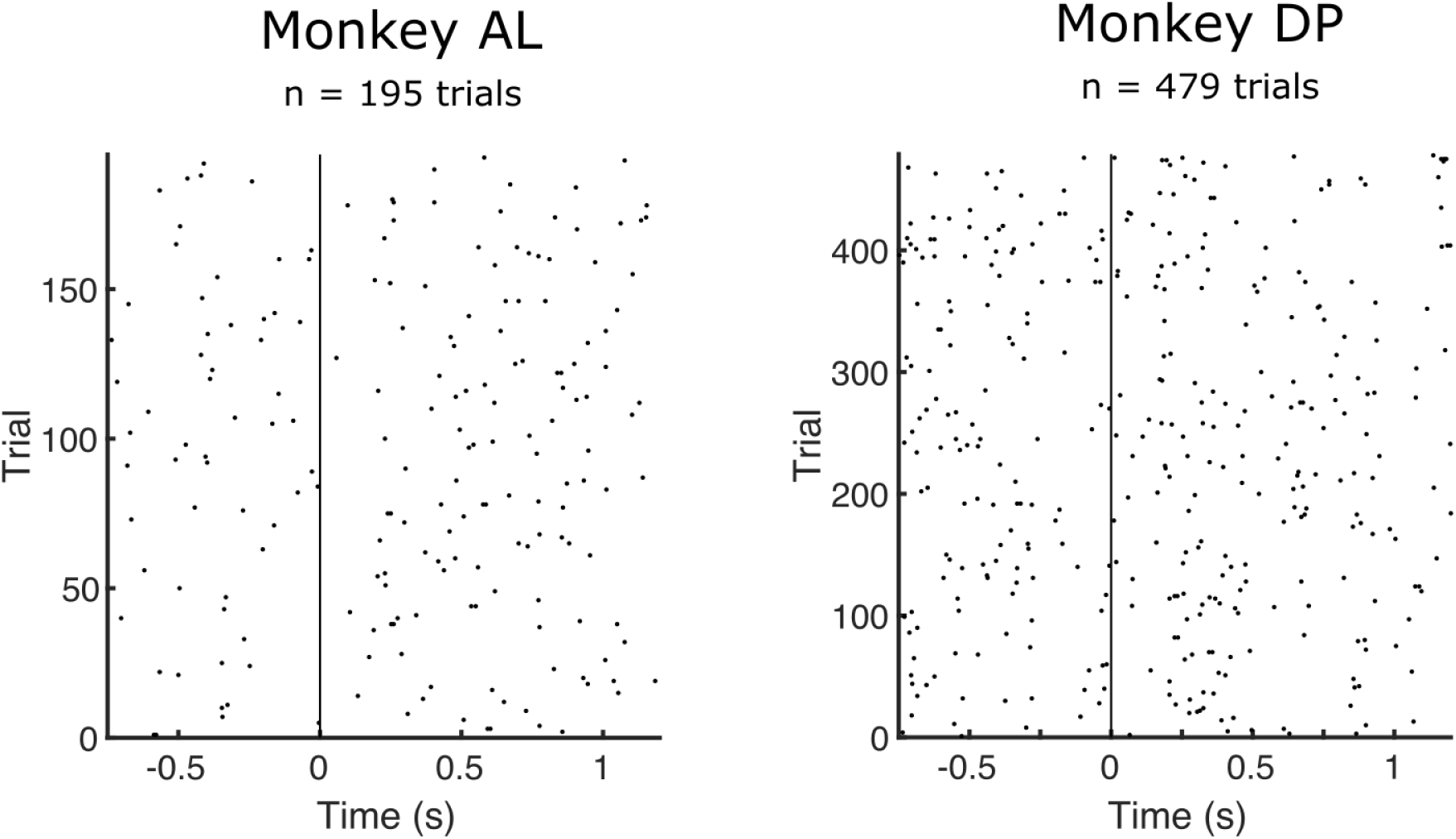
Microsaccade raster plot for size 0.75° combined across sessions. Each dot shows the onset time of microsaccades during a particular trial. The vertical line shows the stimulus onset. There is no apparent rhythmicity in the microsaccades timing across trials.

**Supplementary Figure 8.**
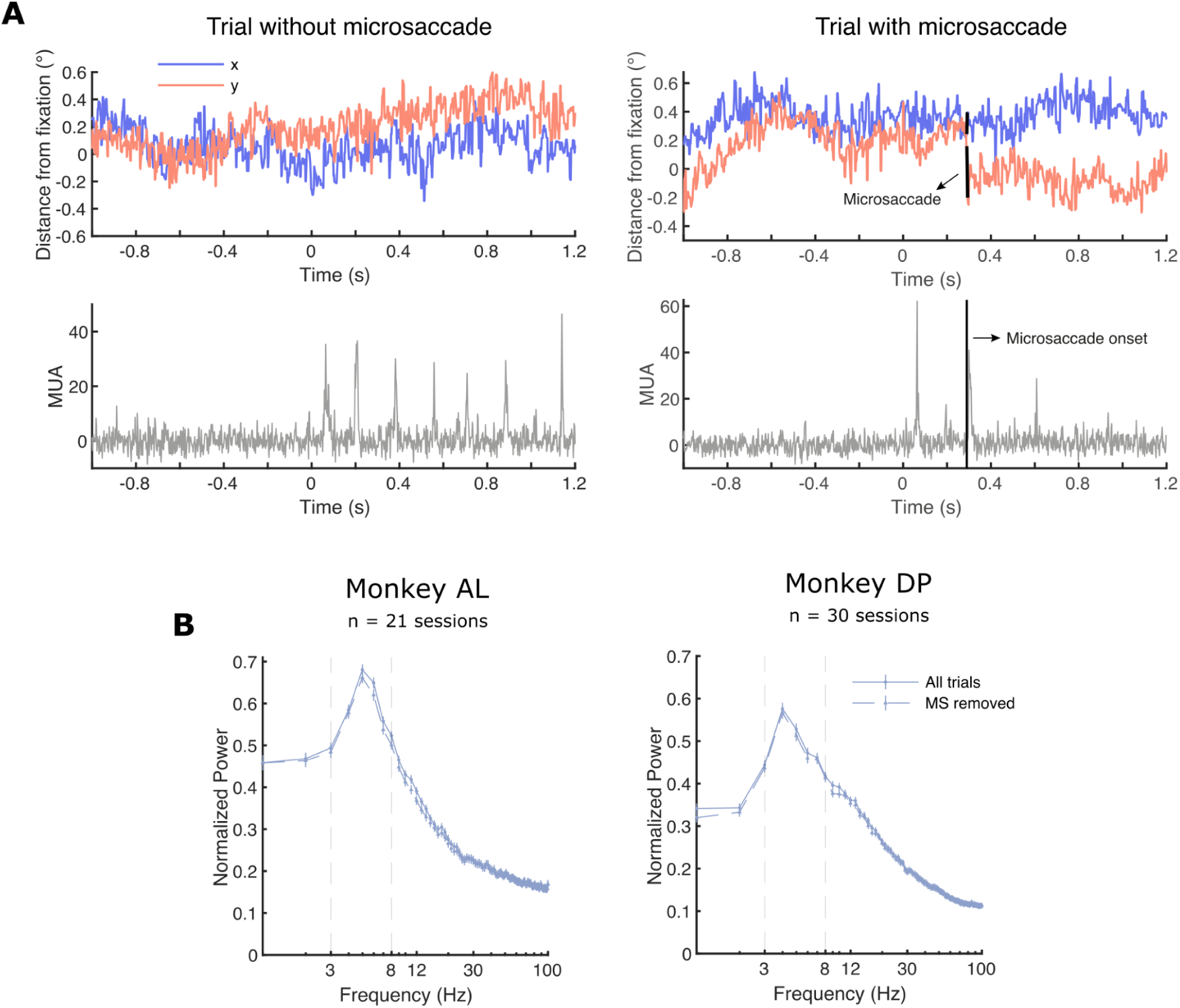
(A) An example trial without (left) and with (right) microsaccade. The top panels show the horizontal (blue) and vertical (red) eye traces. Black lines show a detected microsaccade. The bottom panel shows MUA from the same trials. Notice that on the left panels, theta oscillations are apparent even without a microsaccade. On the right panels, there are no obvious theta oscillations following a microsaccade. (B) Power spectra of stimulus size 0.75° before (solid line) and after (dashed line) removing trials with microsaccades in the post-stimulus period, separately for each monkey. If microsaccades are the generator of neural theta oscillations, we expect their disappearance after the removal of microsaccades. We found that theta oscillations were still present after removing trials with microsaccades.

**Supplementary Figure 9.**
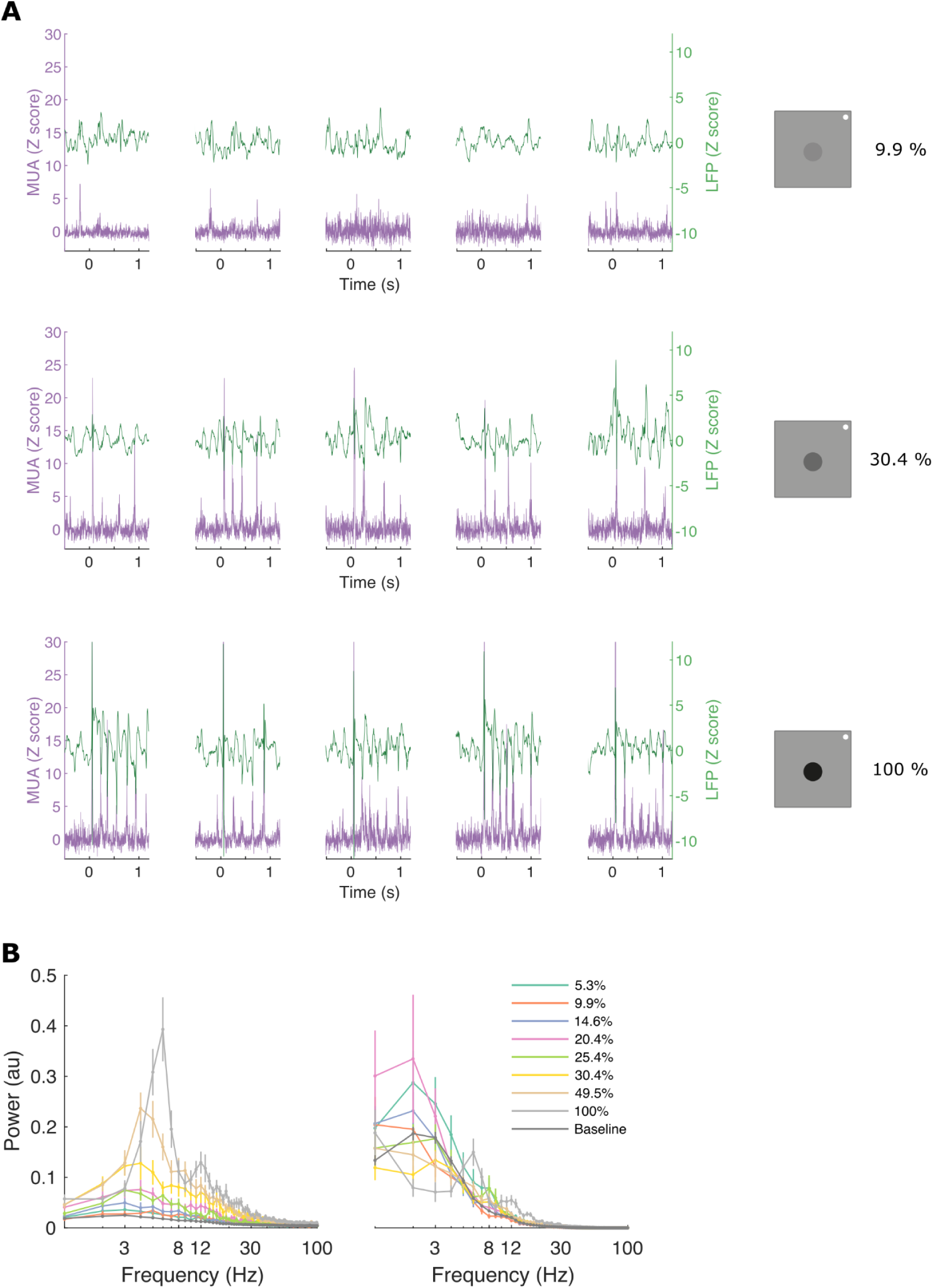
(A) Five example trials of MUA (purple, left y-axis) and LFP (green, right y-axis) response to three different stimulus contrasts on each row Each column corresponds to one trial. Oscillatory activity is clearly seen for 100° stimulus contrast. (B) Mean power spectra of post-stimulus MUA (left) and LFP (right) from the example channel in (A). Dashed lines delineate the theta frequency band.

**Supplementary Figure 10.**
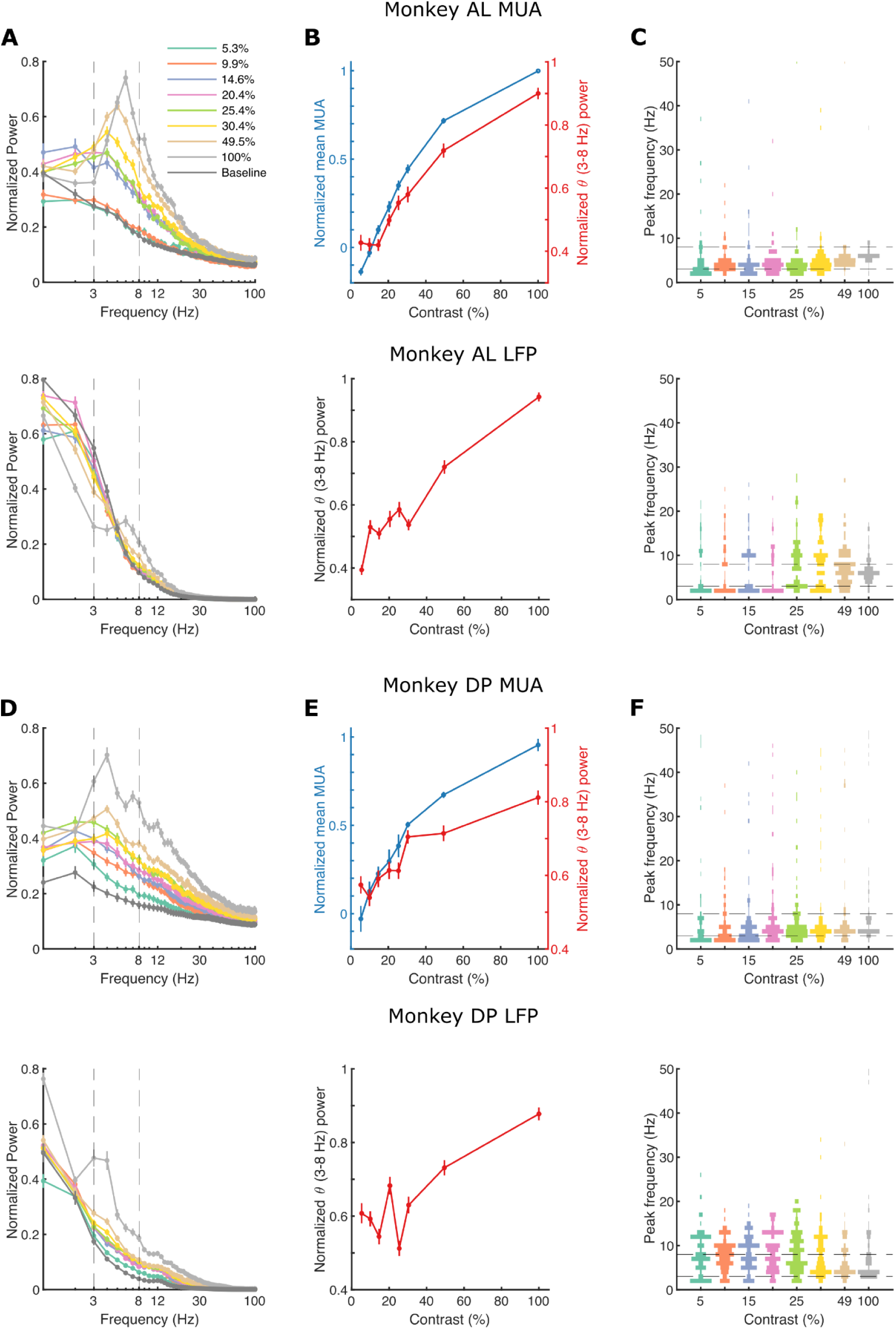
(A) Normalised population MUA (top) and LFP (bottom) power spectra for each contrast, averaged across channels for monkey AL (n = 104 channels). Dashed lines delineate theta frequency range. Error bars are ± 1 standard error of the mean (SEM). (B) Normalised mean MUA (top, blue), MUA theta power (top, red), and LFP theta power (bottom, red) across different contrasts averaged across all channels for monkey AL. Error bars are ± 1 SEM. (C) Peak frequency distribution across channels at different stimulus contrast for monkey AL, for MUA (top) and LFP (bottom) separately. We calculated the mean power spectra across trials for every channel and identified the peak with the strongest power. Theta frequency band is delineated by horizontal dashed lines. (D-F) Same plots but for the monkey DP (n = 126 channels).

## References

Albrecht, D. G., & Hamilton, D. B. (1982). Striate cortex of monkey and cat: Contrast response function. Journal of Neurophysiology, 48(1), 217–237. 10.1152/jn.1982.48.1.217

Allison, J. D., Smith, K. R., & Bonds, A. B. (2001). Temporal-frequency tuning of cross-orientation suppression in the cat striate cortex. Visual Neuroscience, 18(6), 941–948. 10.1017/S0952523801186116

Bosman, C. A., Womelsdorf, T., Desimone, R., & Fries, P. (2009). A Microsaccadic Rhythm Modulates Gamma-Band Synchronization and Behavior. Journal of Neuroscience, 29(30), 9471–9480. 10.1523/JNEUROSCI.1193-09.2009

Busch, N. A., Dubois, J., & VanRullen, R. (2009). The Phase of Ongoing EEG Oscillations Predicts Visual Perception. Journal of Neuroscience, 29(24), 7869–7876. 10.1523/JNEUROSCI.0113-09.2009

Buzsáki, G. (2002). Theta Oscillations in the Hippocampus. Neuron, 33(3), 325–340. 10.1016/S0896-6273(02)00586-X

Carandini, M., Demb, J. B., Mante, V., Tolhurst, D. J., Dan, Y., Olshausen, B. A., Gallant, J. L., & Rust, N. C. (2005). Do we know what the early visual system does? Journal of Neuroscience, 25(46), 10577–10597. 10.1523/JNEUROSCI.3726-05.2005

Cavanaugh, J. R., Bair, W., & Movshon, J. A. (2002). Nature and Interaction of Signals From the Receptive Field Center and Surround in Macaque V1 Neurons. Journal of Neurophysiology, 88(5), 2530–2546. 10.1152/jn.00692.2001

Drebitz, E., Schledde, B., Kreiter, A. K., & Wegener, D. (2019). Optimizing the Yield of Multi-Unit Activity by Including the Entire Spiking Activity. Frontiers in Neuroscience, 13, 83. 10.3389/fnins.2019.00083

Dugué, L., Beck, A.-A., Marque, P., & VanRullen, R. (2019). Contribution of FEF to Attentional Periodicity during Visual Search: A TMS Study. eNeuro, 6(3). 10.1523/ENEURO.0357-18.2019

Dugué, L., Marque, P., & VanRullen, R. (2015). Theta Oscillations Modulate Attentional Search Performance Periodically. Journal of Cognitive Neuroscience, 27(5), 945–958. 10.1162/jocn_a_00755

Dugué, L., Roberts, M., & Carrasco, M. (2016). Attention Reorients Periodically. Current Biology, 26(12), 1595–1601. 10.1016/j.cub.2016.04.046

Engbert, R., & Kliegl, R. (2003). Microsaccades uncover the orientation of covert attention. Vision Research, 43(9), 1035–1045. 10.1016/S0042-6989(03)00084-1

Fiebelkorn, I. C., & Kastner, S. (2021). Spike Timing in the Attention Network Predicts Behavioral Outcome Prior to Target Selection. Neuron, 109(1), 177–188.e4. 10.1016/j.neuron.2020.09.039

Fiebelkorn, I. C., Pinsk, M. A., & Kastner, S. (2018). A Dynamic Interplay within the Frontoparietal Network Underlies Rhythmic Spatial Attention. Neuron, 99(4), 842–853.e8. 10.1016/j.neuron.2018.07.038

Fiebelkorn, I. C., Saalmann, Y. B., & Kastner, S. (2013). Rhythmic Sampling within and between Objects despite Sustained Attention at a Cued Location. Current Biology, 23(24), 2553–2558. 10.1016/j.cub.2013.10.063

Fisher, R. A. (1938). Statistical Methods for Research Workers (7th ed.). Oliver and Boyd.

Foster, K. H., Gaska, J. P., Nagler, M., & Pollen, D. A. (1985). Spatial and temporal frequency selectivity of neurones in visual cortical areas V1 and V2 of the macaque monkey. The Journal of Physiology, 365(1), 331–363. 10.1113/JPHYSIOL.1985.SP015776

Fournier, J., Saleem, A. B., Diamanti, E. M., Wells, M. J., Harris, K. D., & Carandini, M. (2020). Mouse Visual Cortex Is Modulated by Distance Traveled and by Theta Oscillations. Current Biology, 30(19), 3811–3817.e6. 10.1016/j.cub.2020.07.006

Gaillard, C., Ben Hadj Hassen, S., Di Bello, F., Bihan-Poudec, Y., VanRullen, R., & Ben Hamed, S. (2020). Prefrontal attentional saccades explore space rhythmically. Nature Communications, 11(1), Article 1. 10.1038/s41467-020-14649-7

Gao, M., Lim, S., & Chubykin, A. A. (2021). Visual Familiarity Induced 5-Hz Oscillations and Improved Orientation and Direction Selectivities in V1. Journal of Neuroscience, 41(12), 2656–2667. 10.1523/JNEUROSCI.1337-20.2021

Gray, H., Bertrand, H., Mindus, C., Flecknell, P., Rowe, C., & Thiele, A. (2016). Physiological, Behavioral, and Scientific Impact of Different Fluid Control Protocols in the Rhesus Macaque (Macaca mulatta). eNeuro, 3(4). 10.1523/ENEURO.0195-16.2016

Hawken, M. J., Shapley, R. M., & Grosof, D. H. (1996). Temporal-frequency selectivity in monkey visual cortex. Visual Neuroscience, 13(3), 477–492. 10.1017/S0952523800008154

Helfrich, R. F., Fiebelkorn, I. C., Szczepanski, S. M., Lin, J. J., Parvizi, J., Knight, R. T., & Kastner, S. (2018). Neural Mechanisms of Sustained Attention Are Rhythmic. Neuron, 99(4), 854–865.e5. 10.1016/j.neuron.2018.07.032

Jonas. (2025, May 19). Plot spread points (beeswarm plot). https://uk.mathworks.com/matlabcentral/fileexchange/37105-plot-spread-points-beeswarm-plot

Jones, H. E., Grieve, K. L., Wang, W., & Sillito, A. M. (2001). Surround Suppression in Primate V1. Journal of Neurophysiology, 86(4), 2011–2028. 10.1152/jn.2001.86.4.2011

Jutras, M. J., Fries, P., & Buffalo, E. A. (2013). Oscillatory activity in the monkey hippocampus during visual exploration and memory formation. Proceedings of the National Academy of Sciences, 110(32), 13144–13149. 10.1073/pnas.1302351110

Kienitz, R., Cox, M. A., Dougherty, K., Saunders, R. C., Schmiedt, J. T., Leopold, D. A., Maier, A., & Schmid, M. C. (2021). Theta, but Not Gamma Oscillations in Area V4 Depend on Input from Primary Visual Cortex. Current Biology, 31(3), 635–642.e3. 10.1016/j.cub.2020.10.091

Kienitz, R., Schmid, M. C., & Dugué, L. (2022). Rhythmic sampling revisited: Experimental paradigms and neural mechanisms. European Journal of Neuroscience, 55(11–12), 3010–3024. 10.1111/ejn.15489

Kienitz, R., Schmiedt, J. T., Shapcott, K. A., Kouroupaki, K., Saunders, R. C., & Schmid, M. C. (2018). Theta Rhythmic Neuronal Activity and Reaction Times Arising from Cortical Receptive Field Interactions during Distributed Attention. Current Biology, 28(15), 2377–2387.e5. 10.1016/j.cub.2018.05.086

Landau, A. N., & Fries, P. (2012). Attention Samples Stimuli Rhythmically. Current Biology, 22(11), 1000–1004. 10.1016/j.cub.2012.03.054

Lowet, E., Roberts, M. J., Bosman, C. A., Fries, P., & De Weerd, P. (2016). Areas V1 and V2 show microsaccade-related 3-4-Hz covariation in gamma power and frequency. European Journal of Neuroscience, 43(10), 1286–1296. 10.1111/ejn.13126

Michel, R., Dugué, L., & Busch, N. A. (2021). Distinct contributions of alpha and theta rhythms to perceptual and attentional sampling. European Journal of Neuroscience, ejn.15154. 10.1111/ejn.15154

Niell, C. M., & Stryker, M. P. (2010). Modulation of Visual Responses by Behavioral State in Mouse Visual Cortex. Neuron, 65(4), 472–479. 10.1016/j.neuron.2010.01.033

Oostenveld, R., Fries, P., Maris, E., & Schoffelen, J.-M. (2011). FieldTrip: Open Source Software for Advanced Analysis of MEG, EEG, and Invasive Electrophysiological Data. Computational Intelligence and Neuroscience, 2011, 1–9. 10.1155/2011/156869

Ortiz-Rios, M., Haag, M., Balezeau, F., Frey, S., Thiele, A., Murphy, K., & Schmid, M. C. (2018). Improved methods for MRI-compatible implants in nonhuman primates. Journal of Neuroscience Methods, 308, 377–389. 10.1016/j.jneumeth.2018.09.013

Otero-Millan, J., Troncoso, X. G., Macknik, S. L., Serrano-Pedraza, I., & Martinez-Conde, S. (2008). Saccades and microsaccades during visual fixation, exploration, and search: Foundations for a common saccadic generator. Journal of Vision, 8(14), 1–18. 10.1167/8.14.21

Re, D., Inbar, M., Richter, C. G., & Landau, A. N. (2019). Feature-Based Attention Samples Stimuli Rhythmically. Current Biology, 29(4), 693–699.e4. 10.1016/j.cub.2019.01.010

Rollenhagen, J. E., & Olson, C. R. (2005). Low-Frequency Oscillations Arising From Competitive Interactions Between Visual Stimuli in Macaque Inferotemporal Cortex. Journal of Neurophysiology, 94(5), 3368–3387. 10.1152/jn.00158.2005

Siapas, A. G., Lubenov, E. V., & Wilson, M. A. (2005). Prefrontal Phase Locking to Hippocampal Theta Oscillations. Neuron, 46(1), 141–151. 10.1016/j.neuron.2005.02.028

Song, K., Meng, M., Chen, L., Zhou, K., & Luo, H. (2014). Behavioral Oscillations in Attention: Rhythmic α Pulses Mediated through θ Band. Journal of Neuroscience, 34(14), 4837–4844. 10.1523/JNEUROSCI.4856-13.2014

Spyropoulos, G., Bosman, C. A., & Fries, P. (2018). A theta rhythm in macaque visual cortex and its attentional modulation. Proceedings of the National Academy of Sciences, 201719433. 10.1073/pnas.1719433115

Supèr, H., & Roelfsema, P. R. (2005). Chronic multiunit recordings in behaving animals: Advantages and limitations. Progress in Brain Research, 147(SPEC. ISS.), 263–282. 10.1016/S0079-6123(04)47020-4

Tallon-Baudry, C., Bertrand, O., Delpuech, C., & Pernier, J. (1996). Stimulus Specificity of Phase-Locked and Non-Phase-Locked 40 Hz Visual Responses in Human. Journal of Neuroscience, 16(13), 4240–4249. 10.1523/JNEUROSCI.16-13-04240.1996

Vanderwolf, C. H. (1969). Hippocampal electrical activity and voluntary movement in the rat. Electroencephalography and Clinical Neurophysiology, 26(4), 407–418. 10.1016/0013-4694(69)90092-3

VanRullen, R. (2016). How to Evaluate Phase Differences between Trial Groups in Ongoing Electrophysiological Signals. Frontiers in Neuroscience, 10(SEP), 426. 10.3389/fnins.2016.00426

Yoon, S., Baik, B., Park, T., & Nam, D. (2021). Powerful p-value combination methods to detect incomplete association. Scientific Reports, 11(1), 6980. 10.1038/s41598-021-86465-y

Yu, H.-H., Verma, R., Yang, Y., Tibballs, H. A., Lui, L. L., Reser, D. H., & Rosa, M. G. P. (2010). Spatial and temporal frequency tuning in striate cortex: Functional uniformity and specializations related to receptive field eccentricity. European Journal of Neuroscience, 31(6), 1043–1062. 10.1111/j.1460-9568.2010.07118.x

Zimmerman, M. P., Kissinger, S. T., Edens, P., Towers, R. C., Nareddula, S., Nadew, Y. Y., Quinn, C. J., & Chubykin, A. A. (2025). Origin of visual experience-dependent theta oscillations. Current Biology, 35(1), 87–99.e6. 10.1016/j.cub.2024.11.015

Zold, C. L., & Hussain Shuler, M. G. (2015). Theta Oscillations in Visual Cortex Emerge with Experience to Convey Expected Reward Time and Experienced Reward Rate. Journal of Neuroscience. 10.1523/JNEUROSCI.0296-15.2015

